## Supplementary material for "Single molecule microscopy reveals key physical features of repair foci in living cells": Table 1

### List of parameter values

#### WT data

| | |
|---|---|
| Number of cells | 23 |
| Number of traces | 964 |
| Number of displacements | 6252 |
| Average trace length | 7.5 |

| S/G2 Phase | | | |
|---|---|---|---|
| $D_1$ for 2 diffusion population | $1.16 \pm 0.08$ | $\mu m^2/s$ | ML (two-population fit) |
| $D_2$ for 2 diffusion population | $0.28 \pm 0.02$ | $\mu m^2/s$ | ML (two-population fit) |
| Fraction of slow molecules | $0.62 \pm 0.04$ | % | ML (two-population fit) |

| G1 Phase | | | |
|---|---|---|---|
| $D_1$ for 2 diffusion population (from 1 dt) | $1.08 \pm 0.07$ | $\mu m^2/s$ | ML (two-population fit) |
| $D_2$ for 2 diffusion population (from 1 dt) | $0.27 \pm 0.03$ | $\mu m^2/s$ | ML (two-population fit) |
| Fraction of Fast molecules | $0.52 \pm 0.05$ | % | ML (two-population fit) |

#### 1 DSB data

| Quantity | Value | Units | Method |
|---|---|---|---|
| Number of files | 27 | | |
| Number of traces | 1061 | | |
| Number of displacements | 8495 | | |
| Average trace length | 8.0 | | |

| S/G2 Phase | | | |
|---|---|---|---|
| $D_1$ for 3 diffusion population | $1.17 \pm 0.06$ | $\mu m^2/s$ | ML (three-population fit) |
| $D_2$ for 3 diffusion population | $0.24 \pm 0.04$ | $\mu m^2/s$ | ML (three-population fit) |
| $D_3$ for 3 diffusion population | $0.054 \pm 0.007$ | $\mu m^2/s$ | ML (three-population fit) |
| Fraction of D1 molecules | $30 \pm 3$ | % | ML (three-population fit) |
| Fraction of D2 molecules | $38 \pm 4$ | % | ML (three-population fit) |
| Fraction of D3 molecules | $32 \pm 5$ | % | ML (three-population fit) |

| Traces longer than 70 | | | |
|---|---|---|---|
| Confinement radius for molecules | $124 \pm 3$ | $nm$ | From MSD |
| Diffusion Coefficient | $0.032 \pm 0.006$ | $\mu m^2/s$ | From MSD |
| Detection Noise | $28 \pm 2$ | $nm$ | From MSD |

| Cropped data | | | |
|---|---|---|---|
| Confinement radius for molecules | $128 \pm 4$ | $nm$ | From MSD |
| Diffusion Coefficient | $0.038 \pm 0.010$ | $\mu m^2/s$ | From MSD |
| Detection Noise | $30 \pm 3$ | $nm$ | From MSD |

| live PALM | | |
|---|---|---|
| Number of cells | 12 | |
| Radius of Focus | $116 \pm 20$ | nm |
| Number molecules | $880 \pm 600$ | |

#### 2 DSBs data

| Quantity | Value | Units | Method |
|---|---|---|---|
| Number of cells | 14 | | |
| Number of traces | 702 | | |
| Number of displacements | 6220 | | |
| Average trace length | 9.8 | | |
| S/G2 Phase | | | |
| $D_1$ for 3 diffusion population (from 1 dt) | $1.02 \pm 0.05$ | $\mu m^2/s$ | ML (three-population fit) |
| $D_2$ for 3 diffusion population (from 1 dt) | $0.19 \pm 0.02$ | $\mu m^2/s$ | ML (three-population fit) |
| $D_3$ for 3 diffusion population (from 1 dt) | $0.036 \pm 0.004$ | $\mu m^2/s$ | ML (three-population fit) |
| Fraction of D1 molecules | $33 \pm 4$ | % | ML (three-population fit) |
| Fraction of D2 molecules | $48 \pm 4$ | % | ML (three-population fit) |
| Fraction of D3 molecules | $25 \pm 3$ | % | ML (three-population fit) |

| Traces longer than 70 | | | |
|---|---|---|---|
| Confinement radius for molecules | $156 \pm 3$ | $nm$ | From MSD |
| Diffusion Coefficient | $0.041 \pm 0.005$ | $\mu m^2/s$ | From MSD |
| Detection Noise | $28 \pm 3$ | $nm$ | From MSD |

| Cropped data | | | |
|---|---|---|---|
| Confinement radius for molecules | $158 \pm 4$ | $nm$ | From MSD |
| Diffusion Coefficient | $0.040 \pm 0.004$ | $\mu m^2/s$ | From MSD |
| Detection Noise | $30 \pm 2$ | $nm$ | From MSD |

| Live PALM | | |
|---|---|---|
| Number of cells | 11 | |
| Radius of Focus | $170 \pm 20$ | nm |
| Number molecules | $2300 \pm 811$ | |

#### Whole Focus

| | | | |
|---|---|---|---|
| Anomalous exponent | $0.51 \pm 0.05$ | | From MSD |
| Detection Noise | $30 \pm 4$ | $nm$ | From MSD assuming free diffusion in first 8 points |
| Diffusion coefficient | $0.005 \pm 0.002$ | $\mu m^2/s$ | From MSD assuming free diffusion in first 8 points |

#### Rfa1 data

| | | | |
|---|---|---|---|
| Number of cells | 29 | | |
| Number of traces | 621 | | |
| Number of displacements | 7095 | | |
| Average trace length | 12.4 | | |
| Anomalous exponent | $0.56 \pm 0.05$ | | From MSD |
| Detection Noise | $18 \pm 7$ | $nm$ | From MSD assuming free diffusion in first 8 points |
| Diffusion coefficient | $0.0065 \pm 0.0003$ | $\mu m^2/s$ | From MSD assuming free diffusion in first 8 points |
