## Supplementary informations for "Single molecule microscopy reveals key physical features of repair foci in living cells"

Miné-Hattab *et al.*

### SUPPLEMENTARY MATERIAL

#### Functionality test of strains harboring Rad52-halo or Rad52-mMaple

The functionality of strains harboring Rad52-mMaple or Rad52-Halo was tested by dilution assays on plates containing methyl methanesulfonate (MMS). The Rad52-GFP, provided by Mickael Lisby[1,2] is used as a reference strain and exhibits similar viability than the Rad52-mMaple and Rad52-Halo *pdr5Δ* strains used in this study.

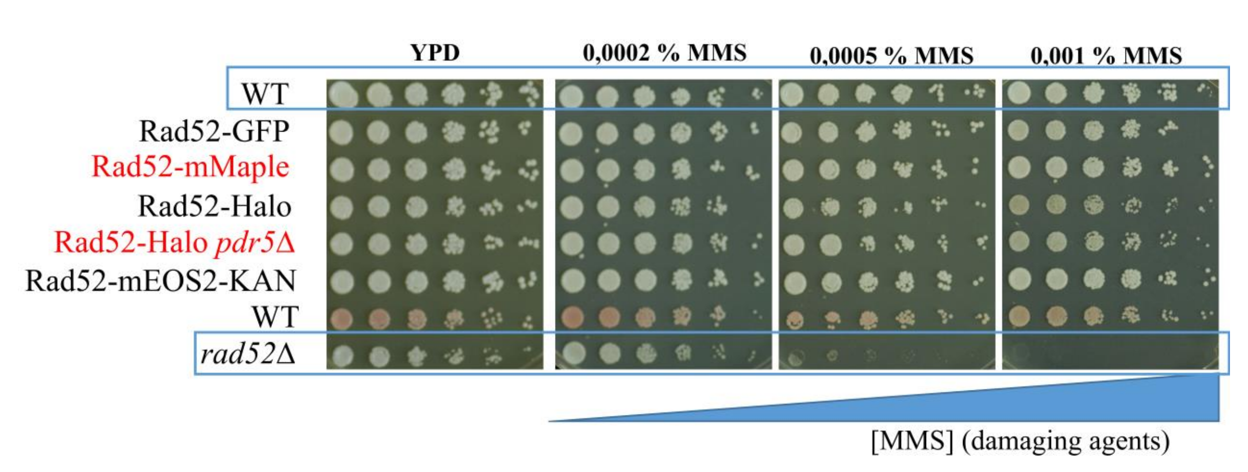


**Figure S1:** Dilution assay of strains harboring Rad52-mMaple or Rad52-Halo on MMS plates.

#### Visualization of Rad52 at the single molecule level

We use a low concentration of JF646 allowing the observation of individual molecules. Figure S2A represents several frames from a typical acquisition: Rad52-Halo-JF646 is visible as a spot in 10 consecutive frames acquired at 50 Hz (20 ms time-intervals). Figure S2B represents the number of detections per frame for a typical acquisition. The top graph corresponds to a nucleus in the absence of DNA damage; the bottom graph represents the number of detection inside a typical Rad52 focus.

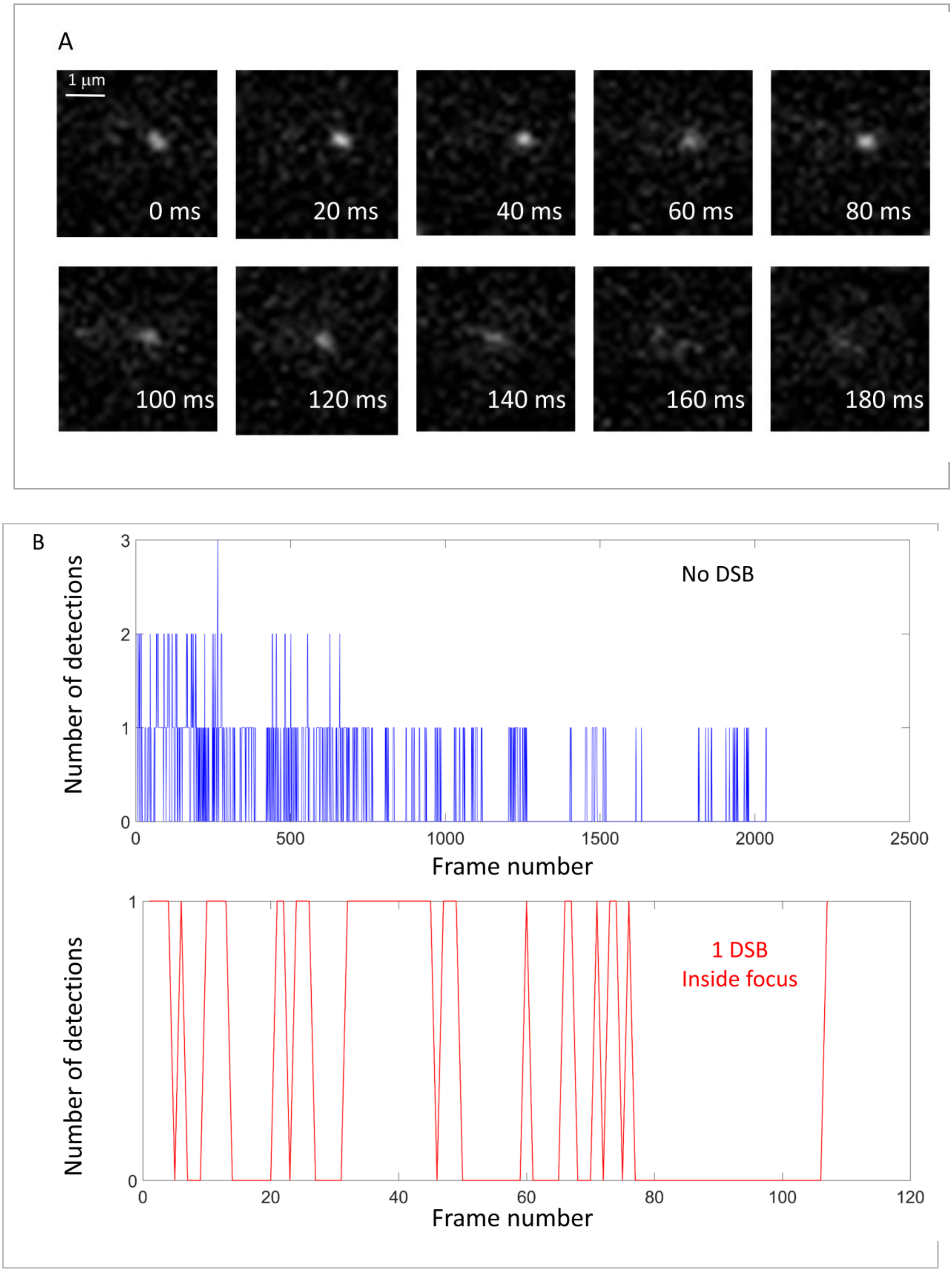


**Figure S2:** (A) Consecutive images from a typical movie in which Rad52-Halo/JF646 are visualized at 20 ms time intervals. (B) Number of Rad52-Halo/JF646 detections per frame for 2 typical cells: Blue: detections per frame inside a cell in the absence of DNA damage induction; Red: detections per frame inside a Rad52 focus in a cell after 2h of I-*Sce*I induction.

**Characterization of JF646 photo-bleaching in budding yeast**

Haploid yeast harboring Rad52-Halo were incubated for 1h with 50 nM of fluorogenic JF646 (see Method) and visualized on a PALM microscope. We then measured the bleaching time of JF646 in 5 cells shown with different colors (insert panel). The average decay (blue curve) was fitted with $Ae^{(-x/\tau)} + cst$, where the half-life of JF646 is estimated at τ = 2.1 seconds.

Of note, the half-life of JF646 is 2.1 s and the longest trace is 1.3 s, the traces length observed in the absence of DSB is due to molecules moving out of focus and not the photo bleaching of the JF646 dyes.

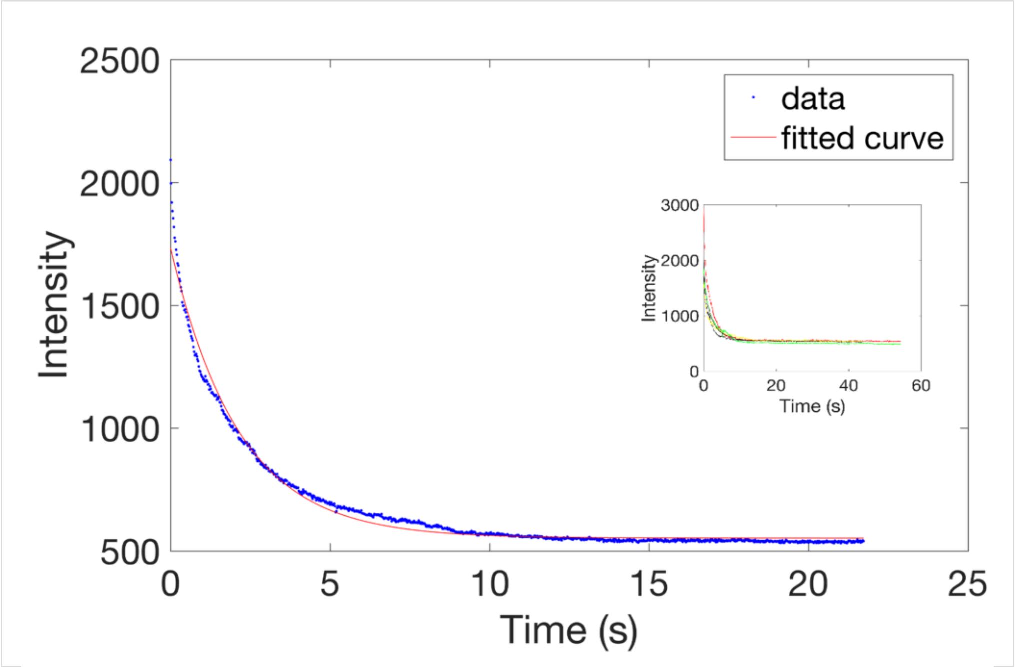


**Figure S3:** JF646 bleaching curve in live budding yeast.

**In the absence of DSB, Rad52 explores the whole nucleus**

In the absence of DSB, Rad52 molecules exhibits confined diffusion (green fitting curve). We obtained a Rad52 confinement radius of  800 nm, close to the radius of haploid yeast nuclei ($R_{haploid}$ = 900 nm, as measured in [3]), indicating that Rad52 explores most of the nucleus in the absence of DSB. We also compared this to an anomalous diffusion fit and found that the confinement fit were in better agreement with the data.

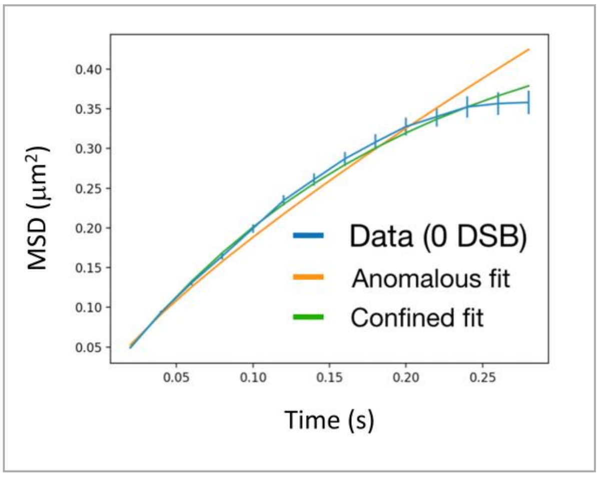


**Figure S4 :** MSD of Rad52-Halo/JF646 in the absence of DSB. The data are fitted with a confined diffusion (green) and an anomalous diffusion (orange).

**Displacement of free Halo-NLS**

We measured the diffusion of free JF as a reference in haploid yeast cells at 20 ms time interval. Since JF646 are fluorogenic, they are visible only when bound to HaloTag. We thus used a haploid strains expressing NLS-Halo. Cells were grown until OD 0.8 and incubated 1h of with 10 nM of JF646.

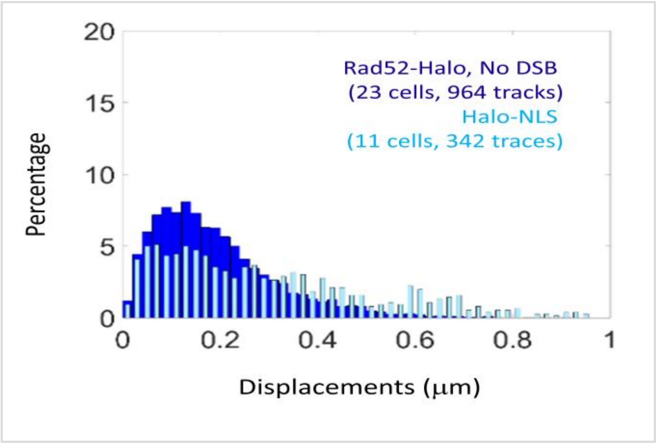


**Figure S5**: light blue: Diffusion of free Halo/JF646 in a S/G2 cells harboring Halo-NLS. Dark blue: diffusion of Rad52-Halo in the absence of DSB (same data as Figure 1E). The time interval is 20 ms for both experiments.

**Criteria to select traces inside repair foci**

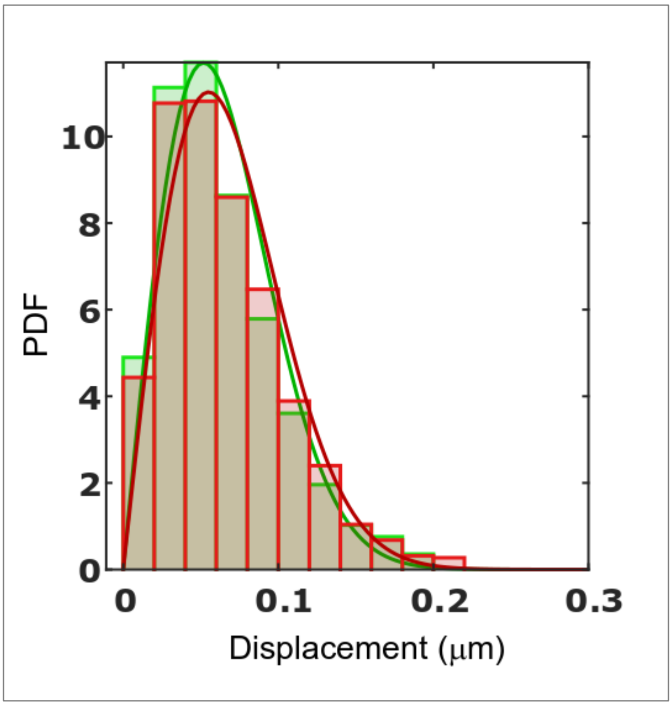


**Figure S6:** PDF of Rad52 traces inside foci induced by a single DSB. Green: selection of traces based on a density threshold; Red: selection of traces based on their length (see Methods). For both method, plain lines represent a 1-population of the data ($p = 0.99$, 2-sided KS test for both methods).

**Nature of motion: whole focus, individual Rfa1 and Rad52 molecules inside foci**

We compared the nature of motion of Rad52 foci and individual Rfa1 and Rad52 molecules inside foci. Using MSD analysis, we observed that Rad52 foci and individual Rfa1 exhibit anomalous motion (Figure S7A and B), whereas Rad52 molecules follow confined motion (Figure S7C). MSD curves are shown in linear scale (Figure S7A, B and C) and in log-log scale (Figure S7D, E and F). The anomalous exponents are 0.51 ± 0.05 and 0.56 ± 0.05 for the whole focus and Rfa1 molecules respectively, as predicted by the Rouse model and consistent with chromatin motion previously measured in the literature[4,5].

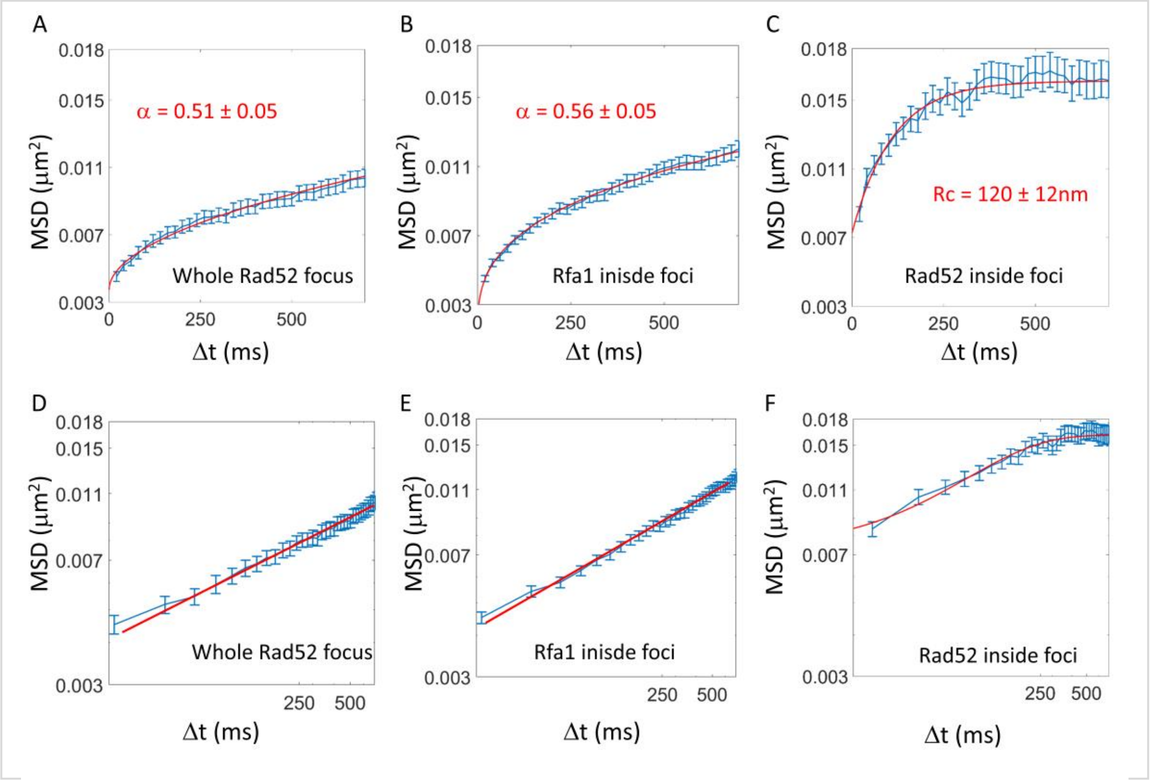


**Figure S7**: (A): Blue: experimental MSD of the whole Rad52 focus in S/G2 cells (14 cells); Red: fit with a model of anomalous diffusion (see Method). (B): Blue: experimental MSD of Rfa1-Halo/JF646 in S/G2 cells (29 cells); Red: fit with a model of anomalous diffusion (see Method). (C) Blue: experimental MSD of Rad52-Halo/JF646 in S/G2 cells (23 cells); Red: fit with a model of confined diffusion (see Method). (D), (E) and (F): same data as (A), (B) and (C) represented in log-log scale. The fits of the MSD are performed on the first 700 ms of the curves.

**Consequences of increasing the concentration of Rad52**

To evaluate the consequences of Rad52 concentration, we measured the concentration of Rad52 in the nucleoplasm for different levels of Rad52 over-expression. We built 2 haploid strains expressing Rad52 under the ADH or TEF1 promoters (5 and 12 times over-expression respectively) [6]. In addition, a single I-*Sce*I cut site allows the induction of a single DSB under the galactose promoter. Cells were grown at 25°C due to a slight growth defect of cells overexpressing Rad52. A single DSB was induced for 2h; during the last hour of induction, cells were incubated with JF646 dyes at 50 nM, a concentration 10 times higher than the one used for single molecule tracking. Such concentration allows the observation of the entire Rad52 focus as a single spot. Cells were observed on an inverted wide field microscope and Rad52 signal was quantified using a home-made software Q-foci[7]. Rad52 concentration is calculated for each nucleus (nucleoplasm intensity divided by nucleoplasm volume).

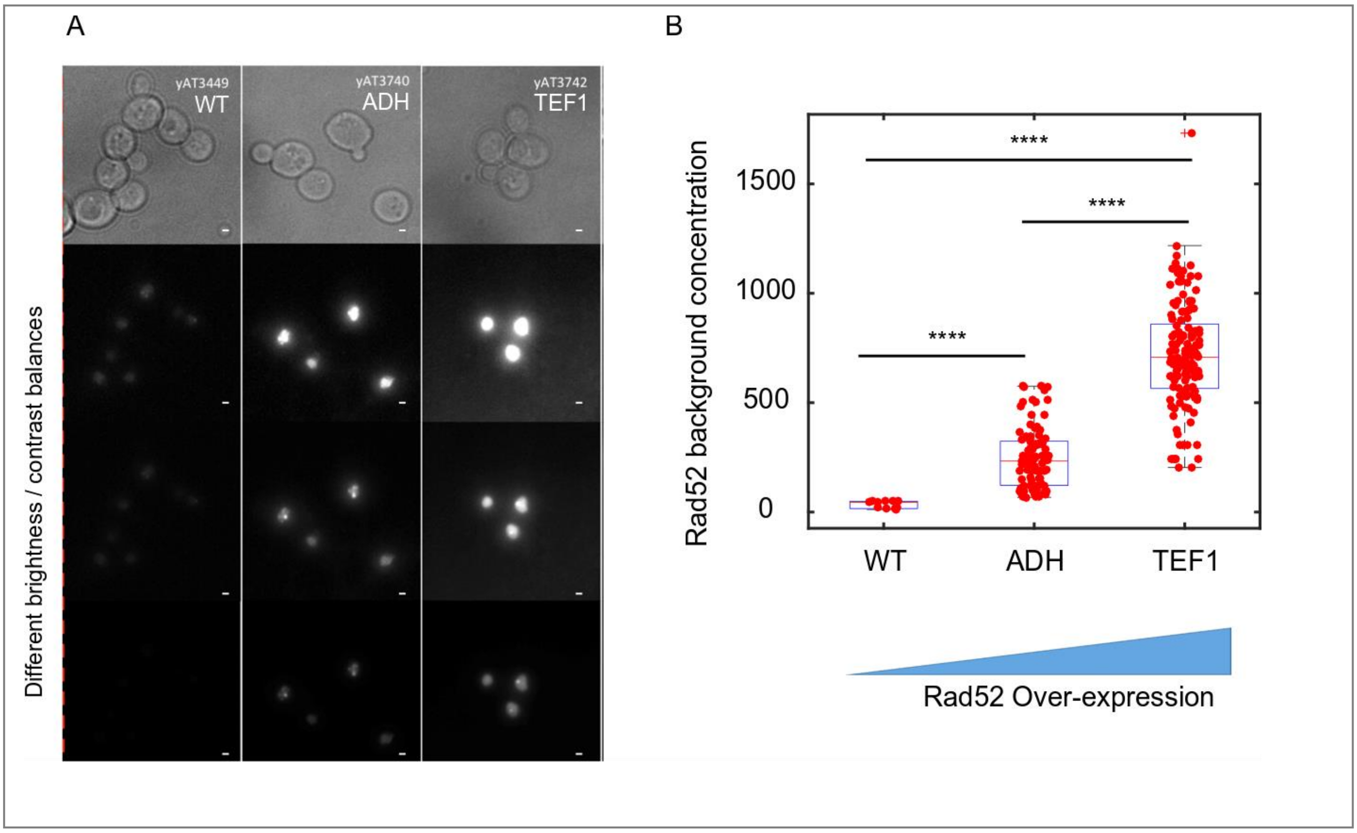


**Figure S8**: (A) Typical images of Rad52 after the induction a single DSB in wild type cells, and for 2 levels of Rad52 over-expression. The bar scale represents 1 μm. (B) Effect of Rad52 over-expression on the nucleoplasm concentration. 11, 94 and 141 cells were analyzed for wild type, ADH and TEF1 promoters respectively.
